# bioPROTACs as versatile modulators of intracellular therapeutic targets: Application to proliferating cell nuclear antigen (PCNA)

**DOI:** 10.1101/728071

**Authors:** Shuhui Lim, Regina Khoo, Khong Ming Peh, Jinkai Teo, Shih Chieh Chang, Simon Ng, Greg L. Beilhartz, Roman A. Melnyk, Charles W. Johannes, Christopher J. Brown, David P. Lane, Brian Henry, Anthony W. Partridge

**Author notes:** **Corresponding Author:** Anthony W. Partridge.

## Abstract

Targeted degradation approaches have recently generated much excitement as a paradigm shift to address human disease in unprecedented ways. Amongst these, small molecule based approaches such as Proteolysis targeting chimeras (PROTACs) have attracted the lion’s share of attention due to their potential to tackle historically intractable targets and achieve greater potency, efficacy, and specificity over traditional small molecule inhibitors. Despite their promise, the identification of high-affinity ligands that can serve as starting points for PROTAC strategies remains challenging. As a complementary approach, we describe herein a class of intracellular biologics termed bioPROTACs. The substrate binding component of these fusion proteins consists of a peptide or an antibody-mimetic which allows for an unprecedented diversity of protein targets that can be addressed. The high-affinity binder is linked directly to an E3 ubiquitin ligase to harness the power of targeted degradation. Using GFP-tagged proteins as model substrates, we show that there is considerable flexibility in both the choice of substrate binders (binding positions, scaffold-class) and the E3 ligases. Indeed, 9 out of 16 binder-E3 combinations tested resulted in greater than 70% target clearance. Through a systematic approach, we then identified a highly effective bioPROTAC against an oncology target, proliferating cell nuclear antigen (PCNA), a sliding DNA clamp with critical roles in DNA replication and repair. The bioPROTAC, termed Con1-SPOP, elicited rapid and robust PCNA degradation and associated effects on DNA synthesis and cell cycle progression. Compared to RNAi-based approaches which typically take days to manifest, PCNA knockdown using Con1-SPOP was evident within 4 h. The advantage of degradation versus stoichiometric inhibition was also clearly demonstrated with bioPROTAC strategies. Combining superior pharmacological inhibition and relative ease of development, bioPROTACs are powerful tools for interrogating the degradability of a substrate, for guiding the identification of the fittest E3 ligase, for studying the functional consequences associated with target protein down-regulation, and potentially for making therapeutic impacts.

## INTRODUCTION

Targeted degradation approaches function by inducing the assembly of the ubiquitination complex in close proximity to a protein-of-interest (POI) to catalyze its selective ubiquitin-tagging and subsequent proteasome-mediated degradation^1^. Several such approaches exist including molecular glues which remodel the surface of an E3 ligase to induce binding to – and degradation of – neo-substrates (e.g. lenolidimide, an approved therapeutic). PROTACs (Proteolysis targeting chimeras), the other major targeted degradation class, are bispecific molecules that induce substrate degradation by simultaneously binding a protein-of-interest (POI) and an E3 ligase (e.g. ARV-110, a degrader of the androgen receptor and the first-in-class PROTAC to enter clinical trials). Pharmacologically, small molecule degraders offer several advantages over traditional inhibitor-based therapeutics. First, degradation can be induced *via* interactions sites across the POI surface, regardless of whether the binding site is of functional consequence^2^, thus expanding the chemical space for tackling otherwise intractable targets^3^. Second, molecules can be recycled for multiple rounds of degradation, a sub-stoichiometric property which is especially useful for high-abundance targets compared to stoichiometric inhibitors may become limited by the high systemic doses required and corresponding polypharmacology-based toxicities^4^. Third, superior pharmacological inhibition can be achieved as degradation attenuates all biological activities (enzymatic, transactivation, scaffolding, etc.) and inhibition is sustained pending protein re-synthesis^5^. Hence, reduced drug dosing frequencies can potentially be realized. Finally, enhanced specificity can be attained through differences in substrate degradability, E3 suitability and ternary complex stability^6–9^. Also, the ability to engage E3s with differences in subcellular localization, cell-cycle dependent regulation or tissue-/disease-specific expression adds an additional layer of selectivity that can be leveraged with targeted degradation strategies.

While small molecule targeted degradation approaches offer compelling advantages, the discovery of corresponding clinical candidates is not without its challenges. For molecular glues, limited examples exist and PROTACs have thus far been only applied to targets with available small molecules inhibitors^10^. Classically ‘undruggable’ proteins remain challenging although opportunities exist to re-purpose previously identified small molecule ligands that did not block protein function for a degradation strategy. Also, out of more than 600 E3 ligases encoded by the human genome, only 4 are routinely used in PROTAC design – CRBN, VHL, MDM2 and cIAP^11^. The choice of the E3 determines degradation efficiencies and the current selection lacks the diversity needed to harness the full potential of the ubiquitin-proteasome system (UPS). Even if small molecule ligands to the POI were available, considerable time and effort (without assurance of success) have to be invested for testing various combinations of linker lengths and E3s recruited. There is currently a poor understanding of the rules that govern stable ternary complex formation between substrate, PROTAC and E3, making informed decisions on PROTAC design difficult. Lastly, as PROTACs are typically composed of 2 ligands connected by a linker, these molecules usually violate Lipinski’s rule of five and thus often suffer from permeability and metabolic liabilities^12^.

As a complementary approach to small molecule based degraders, we sought to develop a biologic equivalent to serve both as a biological tool and as a potential therapeutic approach. Specifically, instead of using small molecules to bridge the substrate and the E3 ligase, we have reengineered the E3 ligase by directly replacing its natural substrate recognition domain with a peptide or a mini-protein that binds a POI. These fusion proteins, which we term bioPROTACs (biological PROTACs), were expressed in cells to drive targeted degradation of POIs. Although bioPROTACs are not novel entities, the work described herein represent a systematic exploration of this approach. Historically, effective degraders have been described for the classically ‘undruggable’ proteins such as β-catenin^13–15^, KRAS^16^ and c-Myc^17^ using domains engrafted from their endogenous interacting partners on to E3 ligases beyond those that can be recruited by small-molecule PROTAC (eg. βTrCP and CHIP). Other proteins also reported to be successfully degraded include cyclin A/CDK2^18^, pRB^19,20^, maltose-binding protein (MBP)^21^, β-galactosidase^21^, and GFP-tagged proteins^22–24^. As a starting point, we further expanded on the published bioPROTAC that degrades GFP-tagged proteins to showcase the remarkable versatility of this system. Building on the insights gained, we used PCNA as an illustrative example to describe and apply a systematic paradigm for the development of potent bioPROTACs against potentially any POI.

## RESULTS

### Validation of vhhGFP4-SPOP for the degradation of H2B-GFP

To establish a model system for identifying active bioPROTAC molecules, we focused on validating published bioPROTACs directed at GFP-tagged proteins since successful turnover can be readily measured through multiple and convenient readouts. In the first report^22^, the region containing the F-box domain from Slmb, a *Drosophila melanogaster* E3, was fused N-terminally to a high-affinity anti-GFP nanobody called vhhGFP4^25,26^. When expressed in various *Drosophila* lines bearing GFP-fusion protein knock-ins, the authors reported effective depletion of GFP signal intensities with the NSlmb-vhhGFP4 chimera. However, we did not achieve knockdown when NSlmb-vhhGFP4 was expressed in mammalian HEK293 cells with stable integration of histone 2B (H2B)-GFP, (**Supplementary Fig. 1**). Since Slmb functions as part of the Cullin-RING E3 ubiquitin ligase (CRL) complex, species-related differences could affect complex formation and ubiquitination efficiencies.

To improve on NSlmb-vhhGFP4, Shin *et al.* swapped NSlmb with other mammalian E3 adaptors from the CRL family and identified a combination, vhhGFP4-SPOP, that successfully degraded H2B-GFP and other GFP-fusion proteins in the nucleus^23^. SPOP (speckle type POZ protein)^27^ is an E3 adaptor protein that functions in complex with cullin-3 (CUL3). The 374-residue protein is comprised of two modular domains, the substrate-binding MATH domain and the CUL3-binding BTB domain separated by a flexible loop. To change the substrate specificity of SPOP and enable the targeting of GFP-tagged proteins, its MATH domain was replaced by vhhGFP4 (Fig. 1a). We adopted the same doxycycline-inducible bidirectional system used in the original publication^23^ to drive co-expression of the bioPROTAC vhhGFP4-SPOP and an mCherry reporter of transfection/expression. After transient transfection and doxycycline induction in HEK293 Tet-On® 3G cells with stable integration of H2B-GFP, GFP and mCherry fluorescence were measured by flow cytometry (Fig. 1b). Consistent with the previous report, GFP intensity was reduced in cells expressing vhhGFP4-SPOP, as marked by the mCherry positive signal. Interestingly, as vhhGFP4-SPOP expression increased, the degradation of H2B-GFP was attenuated, consistent with the well documented ‘hook effect’ seen with small molecule based PROTACS^28^. Control constructs were engineered where either one or both modular components were mutated. Specifically, vhhGFP4_mut_ lacks the complementarity determining region 3 (CDR3) and no longer recognizes GFP whereas SPOP_mut_ lacks the 3-box motif responsible for recruiting CUL3 and thus cannot assemble the ubiquitination machinery. In all controls, H2B-GFP levels were maintained (Fig. 1b). The selective depletion of H2B-GFP in mCherry-positive cells expressing vhhGFP4-SPOP but not its controls was also recapitulated with confocal imaging (Fig. 1c) and Western blot analysis (Fig. 1d). Finally, to demonstrate that H2B-GFP down-regulation was indeed mediated through proteasomal degradation, cells expressing vhhGFP4-SPOP were treated with MG132, a proteasome inhibitor. With increasing concentrations of MG132, the turnover of H2B-GFP was blocked while the levels of FLAG-tagged vhhGFP4-SPOP or mCherry (Fig. 1e and 1f) remained unaffected. This suggested that H2B-GFP was selectively targeted by vhhGFP4-SPOP for degradation in a proteasome-dependent manner.

**Figure 1.**
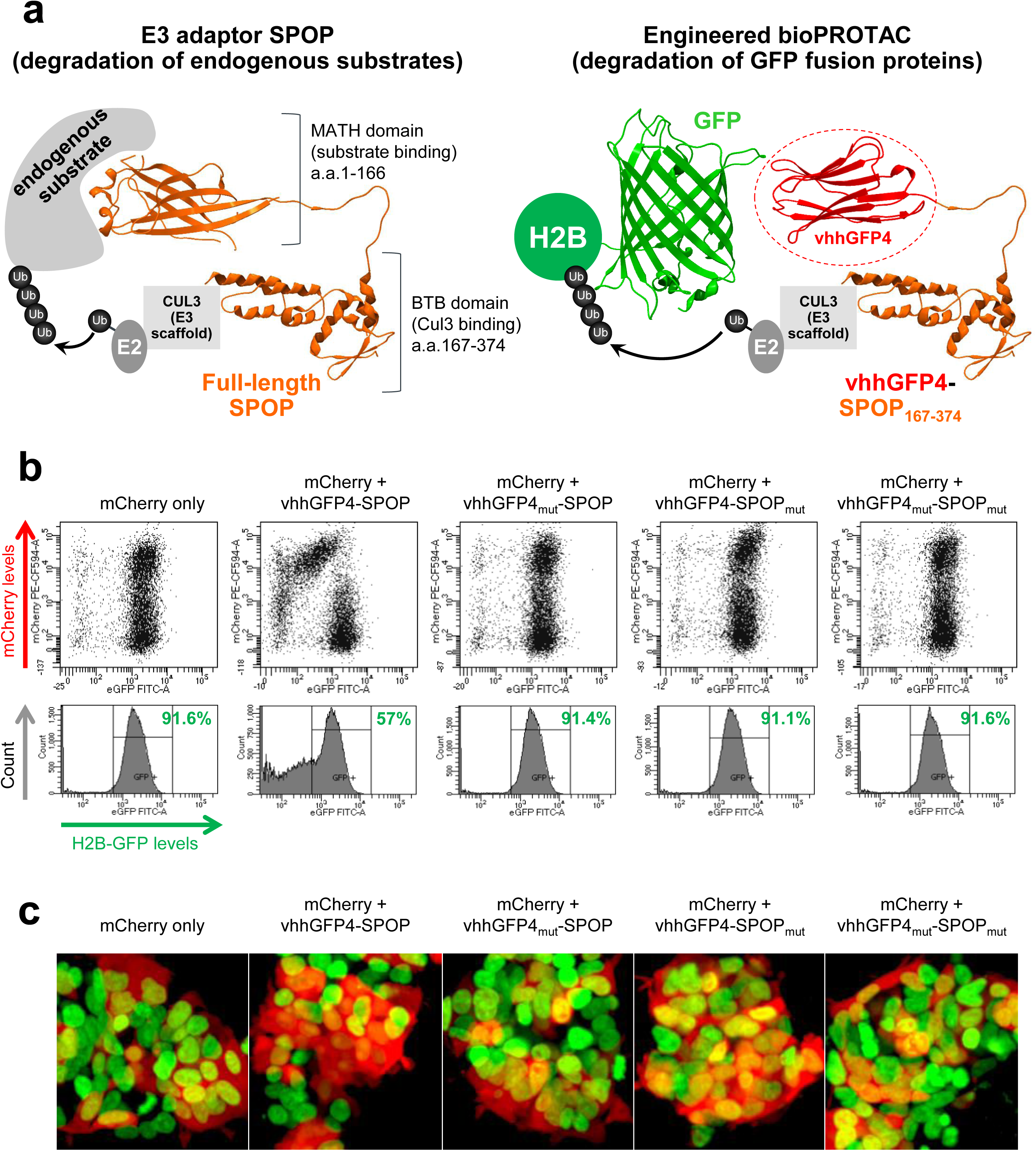
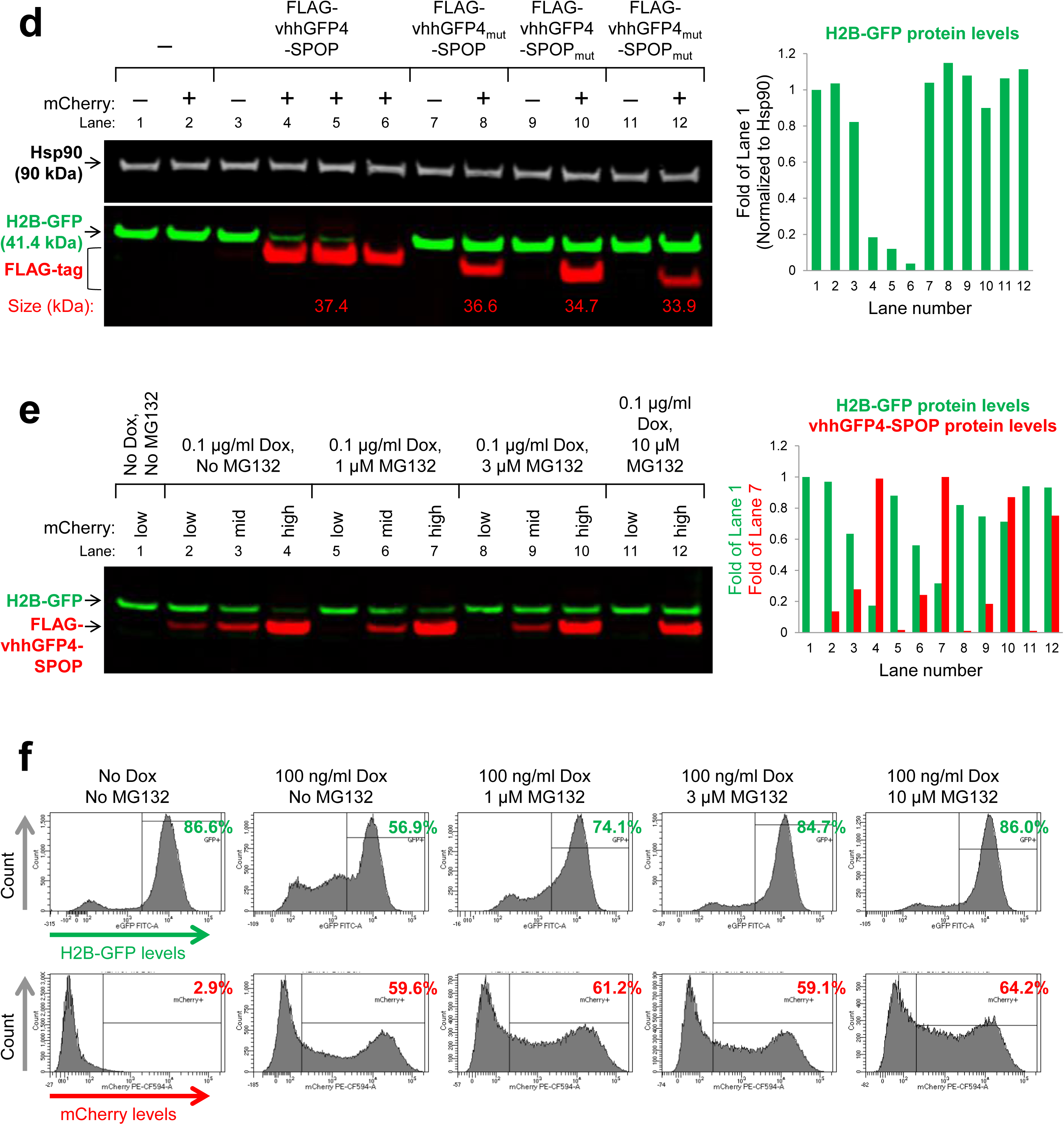
vhhGFP4-SPOP induces the ubiquitin-mediated proteasomal degradation of H2B-GFP. (**a**) Design of the chimeric protein vhhGFP4-SPOP_167-374_ for the degradation of GFP-tagged proteins. The substrate-binding MATH domain of the E3 adaptor SPOP (amino acids 1 to 166) was replaced by vhhGFP4^25,26^, a single-domain antibody fragment that binds GFP. This will enable the ubiquitin-tagging of GFP fusion proteins such as H2B-GFP by the CUL3-based CRL complex. PDB structures are shown for SPOP (3HQI) and GFP:GFP-nanobody complex (3OGO). (**b**) Flow cytometric analysis of H2B-GFP/HEK293 Tet-On® 3G cells transiently transfected with various bidirectional, Tet-responsive plasmids. 100 ng/ml doxycycline was added to induce the simultaneous expression of mCherry and vhhGFP4-SPOP_167-374_ (or its controls). vhhGFP4_mut_ lacks the complementarity determining region 3 (CDR3) and cannot bind GFP. SPOP_mut_ lacks the 3-box motif and cannot bind CUL3. GFP and mCherry fluorescence intensities were measured 24 hours after doxycycline induction. (**c**) Confocal imaging analysis of the same set of cells as in (**b**). (**d**) Western blot analysis of H2B-GFP/HEK293 Tet-On® 3G cells treated as in (**b**) and sorted according to the levels of mCherry using FACS. Gating was set such that mCherry (-) cells have the same signal intensities as untreated cells in the mCherry channel, and anything above this basal level was assigned mCherry (+). Expression of vhhGFP4-SPOP_167-374_ (or its controls) was detected using an anti-FLAG-tag antibody (left panel bottom row, red bands, n=3 for vhhGFP4-SPOP_167-374_) and the expected molecular weight of each chimeric protein is indicated in kilodaltons. The substrate H2B-GFP was detected using an anti-GFP antibody (left panel bottom row, green bands) and band intensities were quantified and normalized to the levels of the loading control Hsp90 (right panel). (**e**) Western blot analysis of H2B-GFP/HEK293 Tet-On® 3G cells transiently transfected with the mCherry/vhhGFP4-SPOP_167-374_ bidirectional inducible plasmid and treated with the indicated concentrations of doxycycline and MG132 (a proteasome inhibitor) for 16 hours. FACS-sorting was conducted as in (**d**). Band intensities of H2B-GFP and FLAG-tagged vhhGFP4-SPOP_167-374_ were quantified and plotted in the right panel. (**f**) Flow cytometric analysis of the same set of cells as in (**e**).

### The extensive flexibility of bioPROTACs

Having validated the PROTAC activity of vhhGFP4-SPOP, we next explored how amenable it was to changes in either the GFP binder or the E3 ligase. From existing literature, we shortlisted a variety of GFP binders that were based on different protein scaffolds, including DARPins^29^, αReps^30^ and monobodies^31^. They were also further diversified by their GFP binding interfaces (Fig. 2a) and reported binding affinities (Fig. 2b). Each was fused to MATH domain deleted SPOP and tested for the ability to degrade H2B-GFP. Surprisingly, despite the drastic differences in size, structure, binding position and affinity, all except the 2 weak binders of GFP (the monobodies GL6 and GL8) were able to deplete GFP signal once expressed (mCherry positive) in HEK293 Tet-On® 3G cells (Fig. 2b). This contrasts with small molecule PROTACs where there was not always a clear correlation between ligand binding affinities and degradation efficiencies. It is important to note that small molecule PROTACs are sandwiched between the substrate and the E3, inducing extensive new protein-protein and protein-ligand contacts that further stabilizes the ternary complex beyond affinities of the individual ligands^6,7^. This does not happen in bioPROTACs and the extent of degradation should be directly proportional to the substrate binding affinity, which helps in simplifying rational design and lead optimization of bioPROTACs.

**Figure 2.**
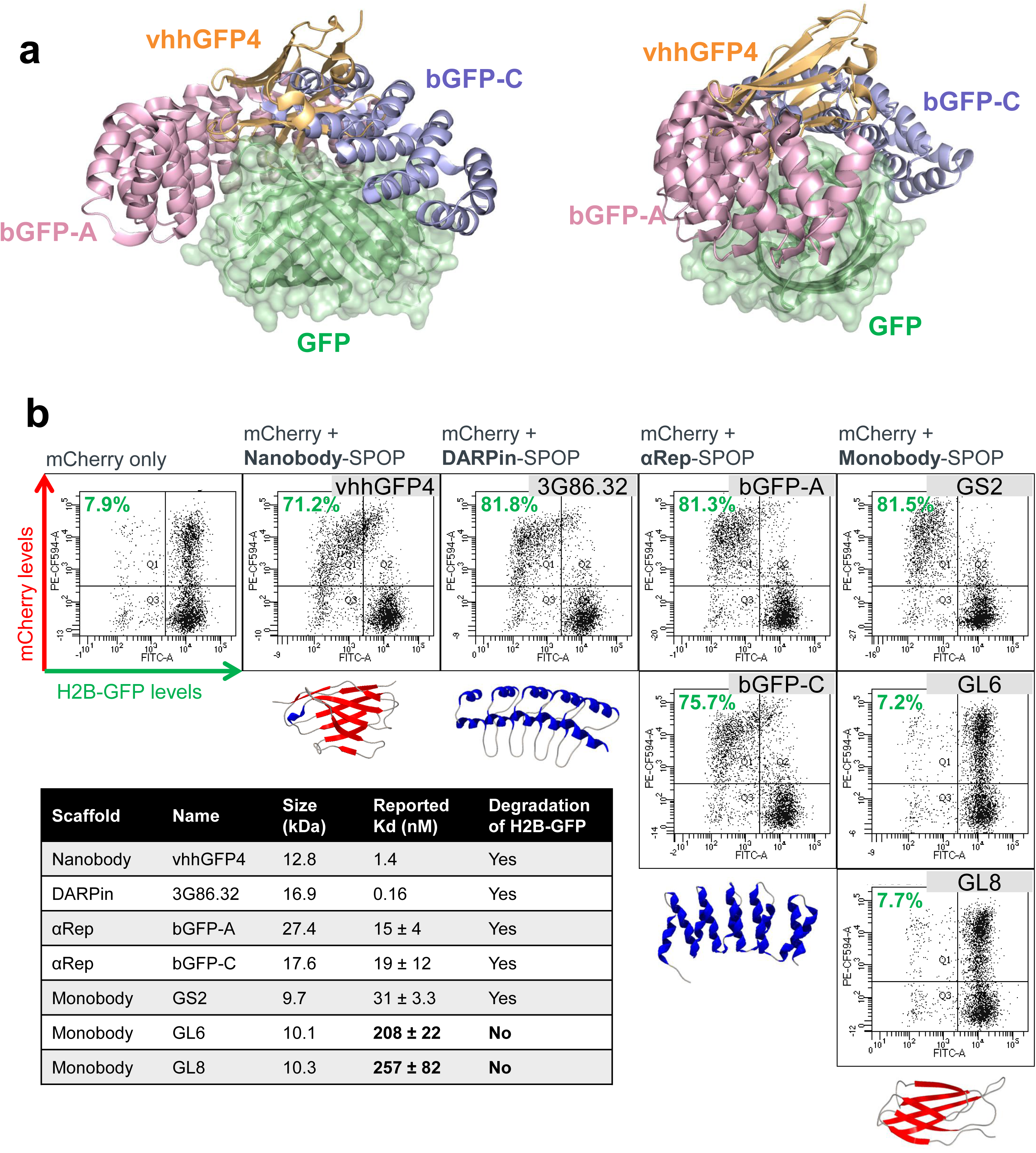
Flexibility in the type of binder used for generating bioPROTACs. (**a**) Overlay of GFP and binders to GFP. 2 different views are shown. vhhGFP4 (PDB 3OGO) is based on the nanobody scaffold while bGFP-A (PDB 4XL5) and bGFP-C (PDB 4XVP) are based on the αRep scaffold. (**b**) Flow cytometric analysis of H2B-GFP/HEK293 Tet-On® 3G cells transiently transfected with various bidirectional, Tet-responsive plasmids. 100 ng/ml doxycycline was added for 24 hours to induce the simultaneous expression of mCherry and the different GFP binders fused to SPOP_167-374_. A total of 7 GFP binders were tested, 1 nanobody (vhhGFP4^25,26^), 1 DARPin (3G86.32^29^), 2 αReps (bGFP-A and bGFP-C^30^) and 3 monobodies (GS2, GL6 and GL8^31^). The values (in green) on the scatter plots indicate the percentage of GFP negative cells in the mCherry positive transfected population, which corresponds to successful H2B-GFP depletion by the respective SPOP-based anti-GFP bioPROTAC. The table summarizes the molecular weight of the GFP binders and their reported binding affinities to GFP. Representative PDB structures are shown for each scaffold (3OGO, 2QYJ, 4XVP, 1TTG), alpha helixes are colored blue and beta strands are colored red.

Through direct fusion of the substrate binder to the E3 ligase, one could potentially recruit any E3-of-interest with a bioPROTAC approach. CRLs are the largest and best-studied family of E3 ubiquitin ligases. They function as multi-subunit complexes that include a cullin scaffold, a RING-H2 finger protein, a receptor responsible for substrate recognition, and with the exception of CUL3-based CRLs, an adaptor subunit that links the substrate receptor to the complex^32^. We picked representative E3 receptors from each of the five major categories of cullins (CUL1 to CUL5). Selection was made based on the availability of structural information to guide truncations. For each E3 receptor, we replaced the substrate recognition domain with vhhGFP4 and retained the portion that binds the remaining E3 complex plus the flexible loop that naturally links the 2 modular domains (Fig. 3a). We also fused vhhGFP4 to the U-box E3 CHIP as it has been proven to be effective in bioPROTAC approaches^16,17,21^. Strikingly, 8 out of 10 of our new bioPROTAC combinations were again successful in degrading H2B-GFP (Fig. 3b), with 5 of them yielding more than 70% clearance. Both CUL4-based CRBN-vhhGFP4 and DDB2-vhhGFP4 failed, suggesting that CUL4 CRLs may be less active in HEK293 cells, or that the protein truncations were not designed optimally resulting in loss of ubiquitin ligase activity. Nevertheless, our results highlighted the versatility of bioPROTACs and the ease at which novel active molecules can be discovered.

**Figure 3.**
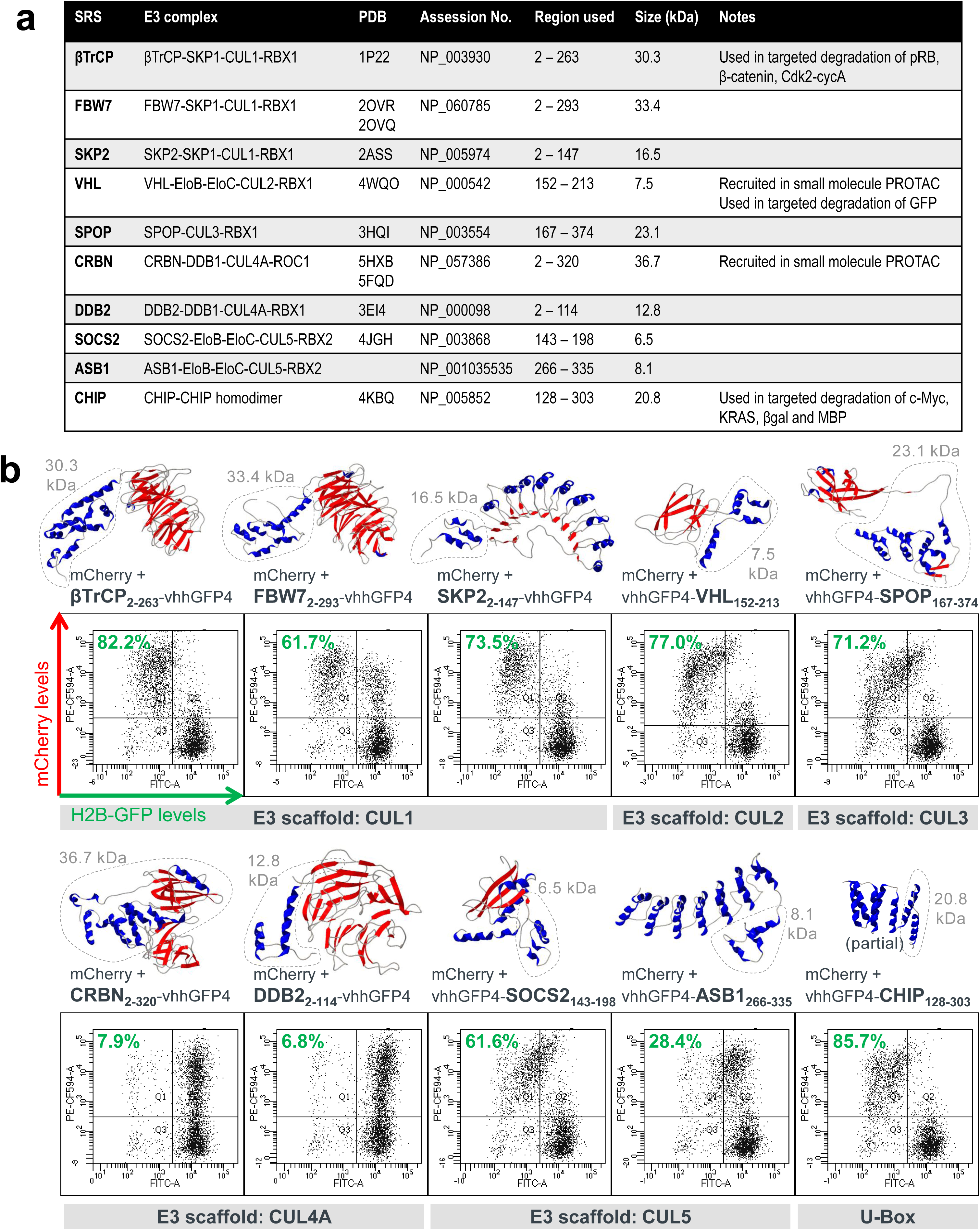
Flexibility in the type of E3 ubiquitin ligase used for generating bioPROTACs. **(a)** Table of 10 different substrate recognition subunit (SRS) used in new bioPROTAC designs to explore alternative E3 ligases. Information regarding each SRS are listed as follows: the E3 ubiquitin ligase complex they function in, PDB structural information, NCBI protein accession number, region of the protein fused to vhhGFP4, molecular weight of the region fused to vhhGFP4 and previous records of their use in PROTAC strategies. Truncations were designed to replace the original substrate-binding domain with the GFP-binding nanobody vhhGFP4. (**b**) Flow cytometric analysis of H2B-GFP/HEK293 Tet-On® 3G cells transiently transfected with various bidirectional, Tet-responsive plasmids. 100 ng/ml doxycycline was added for 24 hours to induce the simultaneous expression of mCherry and the different truncated SRSs fused to vhhGFP4. A total of 10 truncated SRSs were tested, grouped according to the cullin E3 scaffold they recruit. The values (in green) on the scatter plots indicate the percentage of GFP negative cells in the mCherry positive transfected population, which corresponds to successful H2B-GFP depletion by the respective vhhGFP4-based anti-GFP bioPROTAC. PDB structures are shown for each SRS, alpha helixes are colored blue and beta strands are colored red. The dotted area represents the portion fused to vhhGFP4.

### bioPROTACs inform on substrate degradability and E3 selection

The panel of bioPROTACs that degrade GFP-tagged proteins form the basis of our platform to interrogate the degradability of novel substrates as they span representative members in the largest E3 ubiquitin ligase family. For a protein to be degraded through the UPS, a tripartite model has been proposed^33^: 1) a primary degron (short, linear peptide motif) that specifies substrate recognition by cognate E3 ubiquitin ligases, 2) a secondary site comprising of surface lysine(s) that favor (poly)ubiquitin conjugation, and 3) a structurally disordered segment within or proximal to the secondary site such that ubiquitin chain recognition is simultaneously coupled to substrate unfolding at the 26S proteasome. GFP by itself is a poor substrate for ubiquitin-mediated proteasomal degradation (**Supplementary Fig. 2 and Fig. 4a** first row)^22^. Upon the expression of vhhGFP4-SPOP, the localization of GFP switched from uniform cellular distribution to nuclear speckles where SPOP complexes function^34^ (**Supplementary Fig. 2**). However, overall fluorescence intensity was unaffected suggesting that GFP was not efficiently turned over even though it was bound by vhhGFP4-SPOP. Even if GFP was concentrated to the nucleus through the addition of a nuclear localization signal (NLS), the speckles persisted (**Supplementary Fig. 2**). It is unlikely that GFP lacked available lysines for poly-ubiquitination, as 19 lysine residues are evenly distributed across its surface^22^. Instead, due to its compact and well-folded nature, GFP might be missing the flexible sequence that facilitates unfolding and transfer into the catalytic core of the proteasome. When H2B was attached to GFP, GFP signal intensity started to decline upon the expression of various anti-GFP bioPROTACs (**Supplementary Fig. 2 and Fig. 4a** second row), suggesting that the properties which triggered successful proteasomal degradation were imparted by H2B. Therefore, by tagging any POI to GFP, we are now able to recruit a representative pool of E3 ligases to evaluate if the POI possesses the necessary traits that enable its targeted proteolysis – ubiquitin-acceptor surface lysine(s) located within or proximal to structurally disordered degradation initiation sites^33^.

To extend our work to a potential therapeutic target, we sought to determine if PCNA is a good substrate for ubiquitin-mediated proteasomal degradation. PCNA is an essential protein expressed in the nuclei of all proliferating cells where it forms a homo-trimeric ring structure encircling DNA^35^. PCNA serves as a sliding DNA clamp to recruit a myriad of DNA replication and damage repair proteins to the chromatin. Expression of PCNA is elevated in rapidly dividing tumor cells and in most cases is associated with poor prognosis, making it an attractive target for cancer therapy^36^. Considering its scaffolding function and the multiple protein-protein and protein-DNA interactions that PCNA is involved with, PROTAC strategies are ideally poised to abolish all its activities concurrently and achieve therapeutic efficacy. Prior to our study, it was not clear if PCNA levels could be modulated through proteolysis, although numerous reports have described the extensive regulatory mono- and poly-ubiquitination of PCNA in response to genotoxic stress^37,38^. These non-proteolytic signaling events help to coordinate the dynamic engagement of PCNA with its vast array of interacting partners. Since surface lysines are available for ubiquitin conjugation, the appropriate E3s could be recruited to extend the right linkages that target PCNA to the proteasome. To test this hypothesis, we tagged PCNA with GFP and applied our panel of anti-GFP bioPROTACs. 5 out of the 10 tested – vhhGFP4 fused to FBW7 (CUL1), VHL (CUL2), SPOP (CUL3), SOCS2 (CUL5) and CHIP (U-box), were able to deplete PCNA-GFP (Fig. 4a third row). Since the GFP-tag on its own was not efficiently degraded, this result indicated that PCNA possesses the necessary traits that enabled its targeted degradation. Amongst the active E3 ligases identified, SPOP was of specific interest as it shares the same subcellular localization as PCNA – the nucleus. Indeed, the silencing of PCNA-GFP by vhhGFP4-SPOP but not its controls was corroborated with imaging (Fig. 4b). More importantly, SPOP-based bioPROTACs displayed immense flexibility as swapping the nanobody vhhGFP4 with either the DARPin 3G86.32, the monobody GS2, or the αReps bGFP-A/C all generated effective bioPROTACs that downregulated PCNA-GFP (Fig. 4b). Hence, by tagging novel POIs such as PCNA to GFP and applying the anti-GFP bioPROTAC platform that we have built, we are able to probe if the POI can be targeted for proteasomal degradation, and also to identify suitable E3 ligases to recruit for its polyubiquitination.

**Figure 4.**
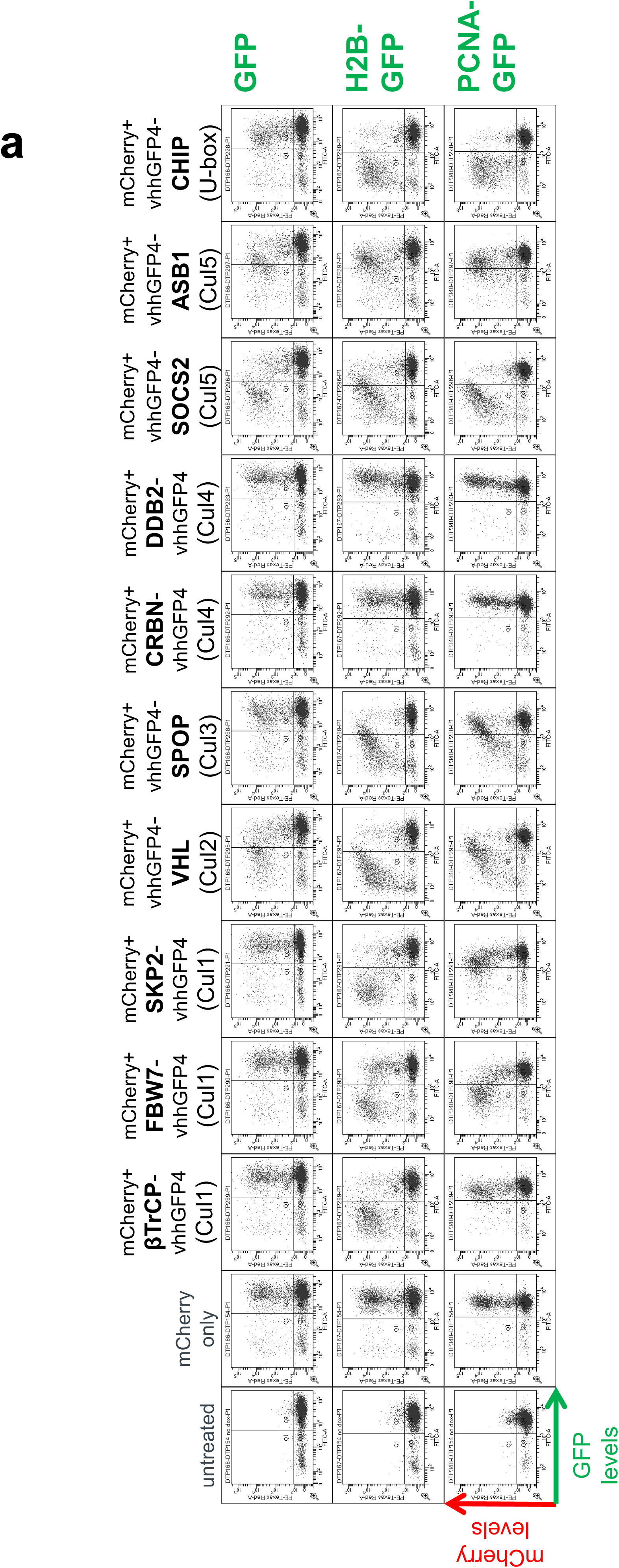
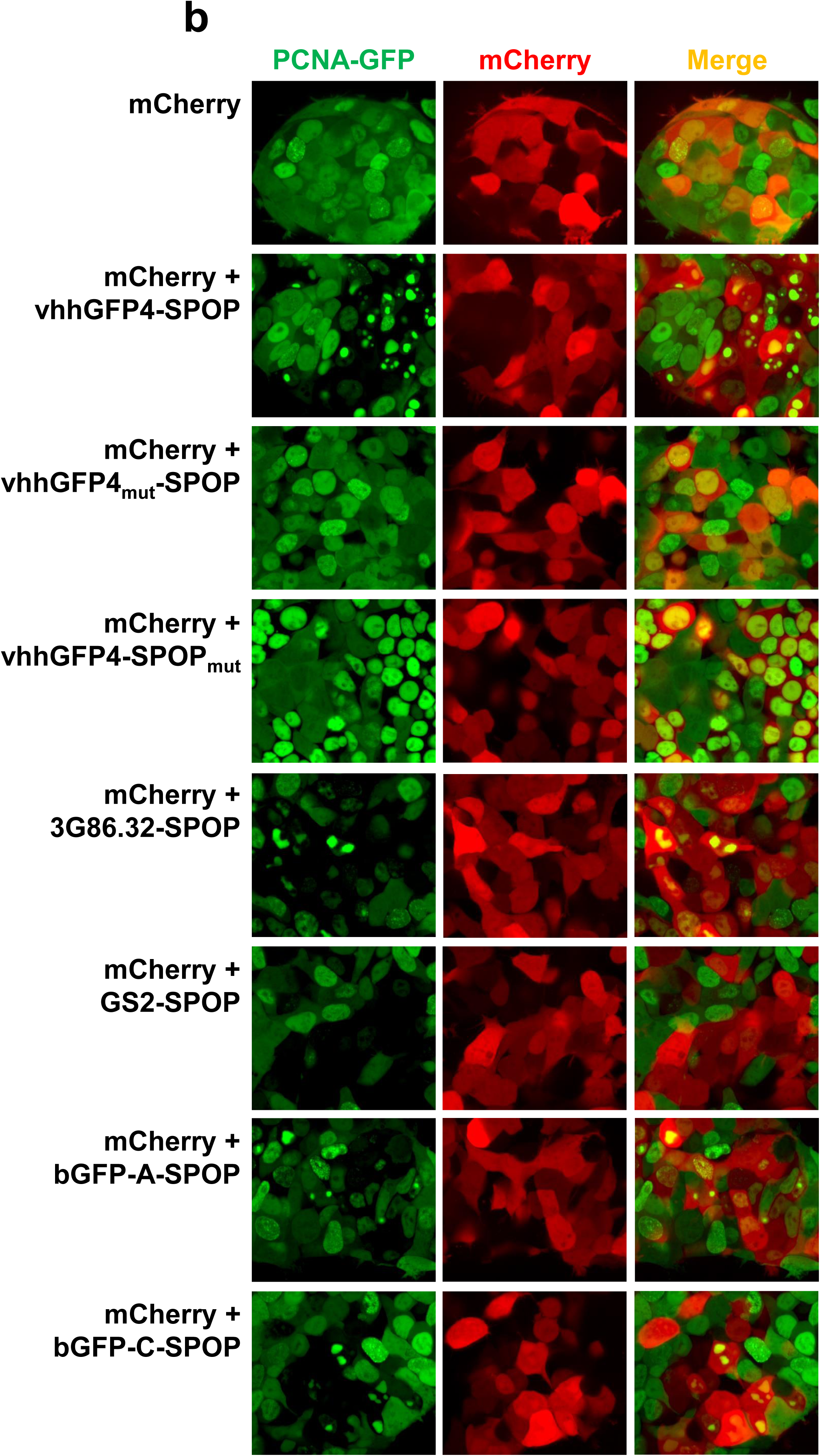
bioPROTACs inform on substrate degradability and E3 selection. (**a**) Flow cytometric analysis of HEK293 Tet-On® 3G cells with stable integration of GFP, H2B-GFP or PCNA-GFP. Each of the 3 stable cell lines was transiently transfected with the same panel of bidirectional, Tet-responsive plasmids. 100 ng/ml doxycycline was added for 24 hours to induce the simultaneous expression of mCherry and the different anti-GFP bioPROTAC. Cells in Q1 represent successful H2B-GFP depletion by the respective anti-GFP bioPROTAC. (**b**) Confocal imaging analysis of PCNA-GFP/HEK293 Tet-On® 3G cells transiently transfected with various bidirectional, Tet-responsive plasmids. 100 ng/ml doxycycline was added for 24 hours to induce the simultaneous expression of mCherry and the different anti-GFP bioPROTACs.

### Con1-SPOP induced robust degradation of PCNA

To verify if the findings from the anti-GFP bioPROTAC platform can be translated to the degradation of endogenous PCNA, we selected a published PCNA-binding peptide termed Con1 to be incorporated into our anti-PCNA bioPROTAC design (Fig. 5a). The 16-residue Con1 peptide binds PCNA with a reported Kd of 100 nM and contains the conserved PIP (PCNA-interacting protein) box motif common to PCNA binding partners^39^. Con1 was fused to SPOP_167-374_ (Fig. 5a), the E3 ligase identified through our anti-GFP bioPROTAC screen. Similar to vhhGFP4-SPOP (with specificity for GFP), Con1-SPOP (with specificity for PCNA) also degraded PCNA-GFP, and furthermore appeared to do so more effectively (53.4% versus 85.8% GFP-negative cells at 24 h post-expression, Fig. 5b). By replacing the 3 conserved residues critical for binding to PCNA in Con1 by alanine, Con1_mut_-SPOP was no longer able to bind and degrade PCNA-GFP. By deleting the 3-box motif responsible for recruiting CUL3, Con1-SPOP_mut_ also failed to alter the levels of PCNA-GFP suggesting that proper assembly of the ubiquitination complex was needed to drive the down-regulation of PCNA-GFP (Fig. 5b).

**Figure 5.**
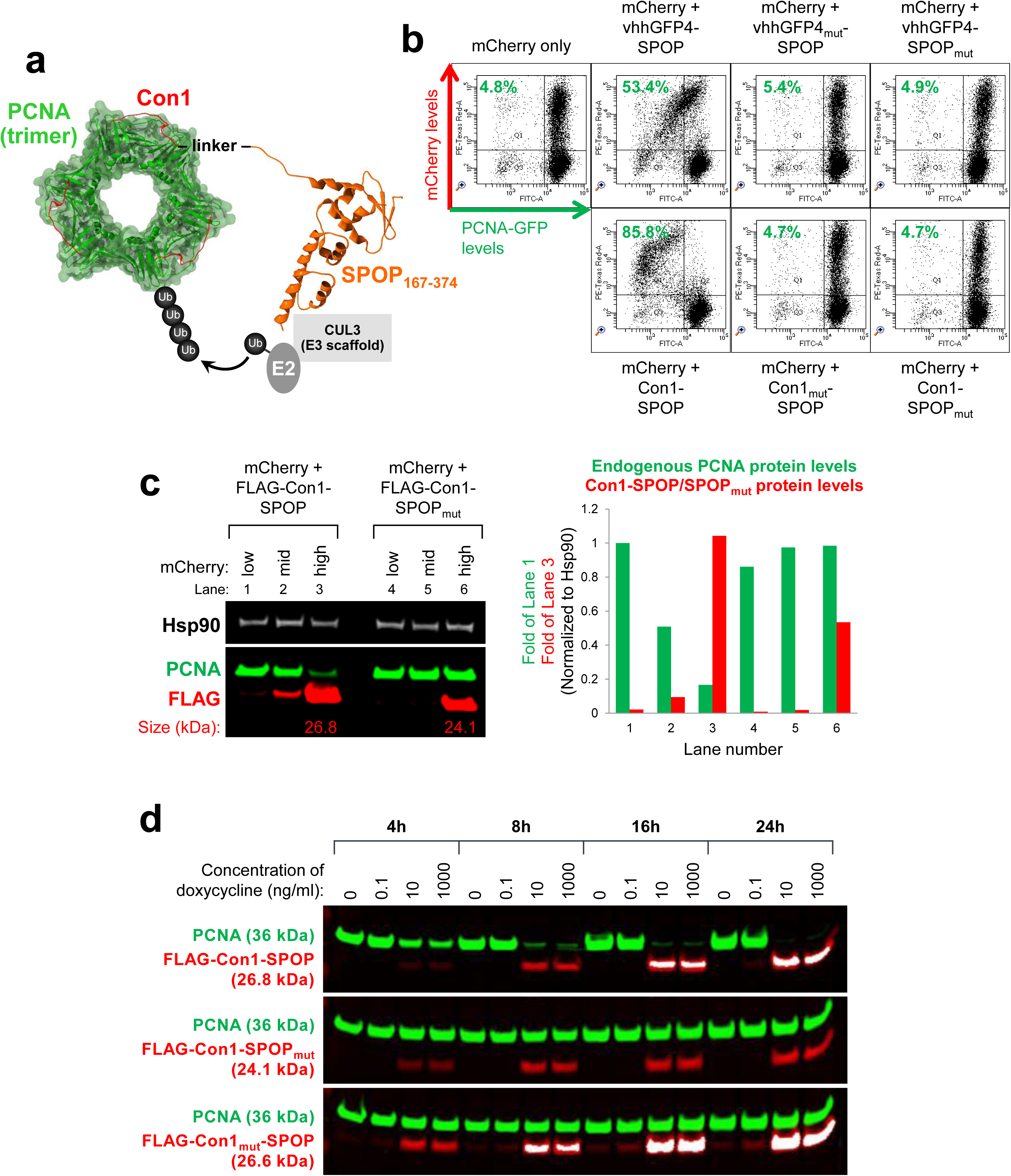
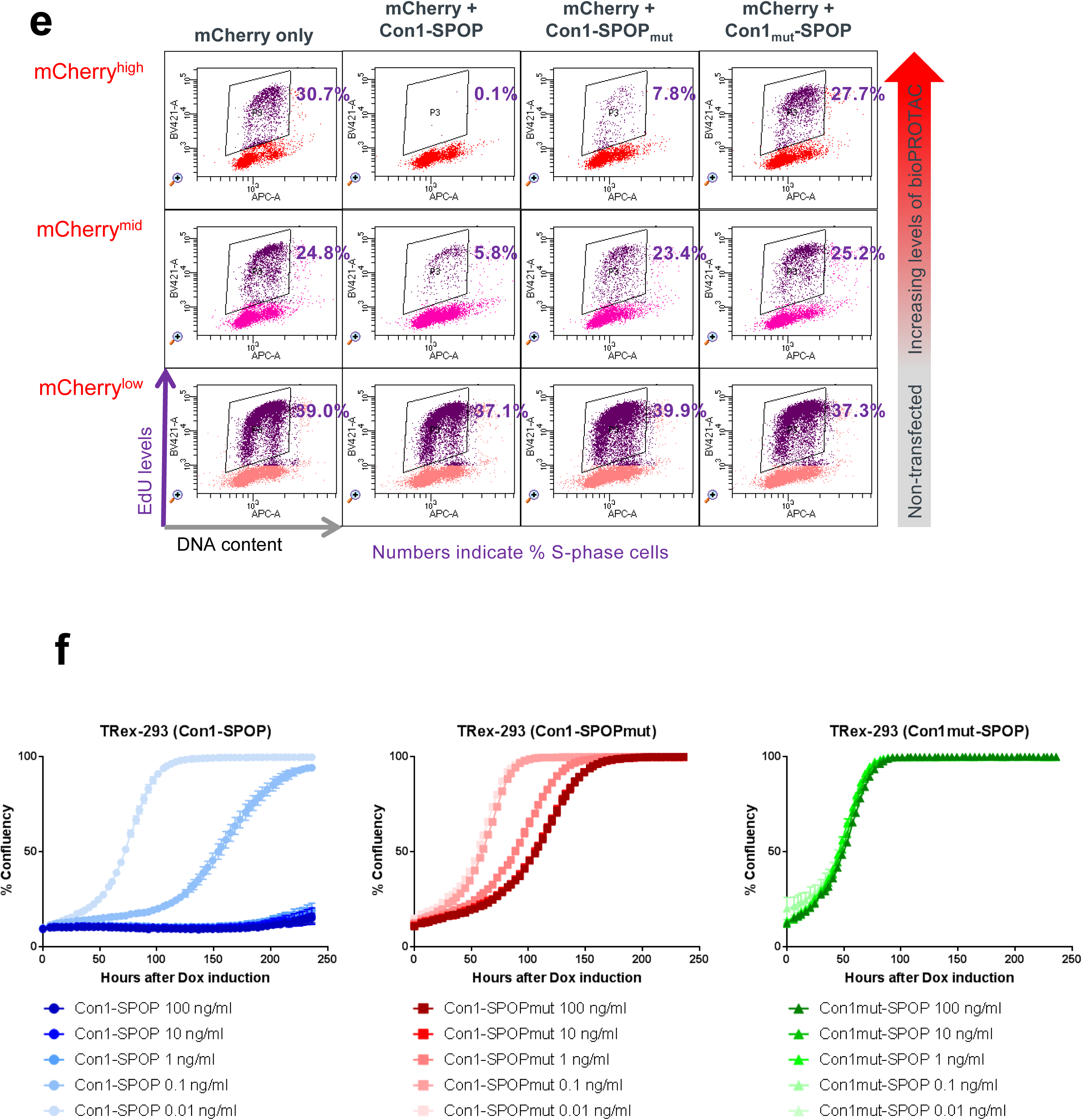
Rapid and robust degradation of PCNA with Con1-SPOP. (**a**) Design of the chimeric protein Con1-SPOP_167-374_ for the degradation of PCNA. The substrate-binding MATH domain of the E3 adaptor SPOP (amino acids 1 to 166) was replaced by Con1, a high affinity peptide ligand of PCNA. This will enable the ubiquitin-tagging of PCNA by the CUL3-based CRL complex. PDB structures are shown for SPOP (3HQI) and PCNA (1AXC). **(b)** Flow cytometric analysis of PCNA-GFP/HEK293 Tet-On® 3G cells transiently transfected with various bidirectional, Tet-responsive plasmids. 100 ng/ml doxycycline was added for 24 hours to induce the simultaneous expression of mCherry and the different chimeric proteins. vhhGFP4_mut_ lacks the complementarity determining region 3 (CDR3) and cannot bind GFP. SPOP_mut_ lacks the 3-box motif and cannot bind CUL3. Con1_mut_ bears point-mutations in the 3 critical PCNA-interacting residues and cannot bind PCNA. The values (in green) on the scatter plots indicate the percentage of GFP negative cells in the mCherry positive transfected population, which corresponds to successful PCNA-GFP depletion by the respective SPOP-based bioPROTAC. (**c**) Western blot analysis of HEK293 Tet-On® 3G cells transiently transfected and induced with doxycycline as in (**b**) and sorted according to the levels of mCherry using FACS. Gating was set such that mCherry^low^ cells have the same signal intensities as untreated cells in the mCherry channel, whereas mCherry^mid^ and mCherry^high^ cells have increasing levels of mCherry fluorescence. Expression of Con1-SPOP_167-374_ (or its control) was detected using an anti-FLAG-tag antibody (left panel bottom row, red bands) and the expected molecular weight of each chimeric protein is indicated in kilodaltons. The substrate PCNA was detected using an anti-PCNA antibody (left panel bottom row, green bands). Band intensities of FLAG-tagged vhhGFP4-SPOP_167-374_/SPOP_mut_ and endogenous PCNA were quantified and normalized to the levels of the loading control Hsp90 (right panel). (**d**) Western blot analysis of T-REx™-293 cells with stable integration of Con1-SPOP_167-374_ (or its controls) under the control of a Tet-responsive promoter. Various concentrations of doxycycline were added to the culture media for the indicated length of time and lysates were collected. Proteins were detected as in (**c**). (**e**) EdU labeling and flow cytometric analysis of HEK293 Tet-On® 3G cells transiently transfected and induced with doxycycline as in (**b**). Cells undergoing DNA synthesis were labeled with 10 µM EdU for 2 hours and the percentage of EdU-positive S-phase cells (in purple) was expressed according to the level of mCherry signal intensities. Gating for mCherry expression was performed as in (**c**). (**f**) Incucyte confluency measurements of T-REx™-293 cells with stable integration of Con1-SPOP_167-374_ (or its controls) under the control of a Tet-responsive promoter. Various concentrations of doxycycline were added to the culture media and the percentage confluency of the cells was tracked continuously over 10 days.

We next investigated the ability of Con1-SPOP to degrade endogenous PCNA. In HEK293 cells, as the expression of FLAG-tagged Con1-SPOP increases (inferred by increases in mCherry signal), the protein levels of PCNA dropped correspondingly (Fig. 5c). In the control where Con1 is able to bind PCNA but does not degrade it, PCNA levels were maintained as expected (Fig. 5c). Using a doxycycline-inducible line with stable integration of Con1-SPOP, PCNA down-regulation was observed as early as 4 hours following the addition of doxycycline, and by 24 hours PCNA protein was barely detectable (Fig. 5d). This effect was again lost when either SPOP or Con1 were mutated (Fig. 5d). The rapid silencing of PCNA demonstrated in this study using bioPROTACs contrasts with previous studies using siRNAs, where more than 72 hours was needed to achieve knock-down^40–43^-an observation that is in-line with PCNA’s reported protein half-life of 78.5 hours^44^. Indeed, a key distinguishing feature of targeted degradation approaches is that the protein target is directly depleted whereas RNA-interference approaches depend on the natural turnover of the existing pool of proteins while preventing *de novo* protein synthesis. In this study, we present a novel bioPROTAC approach to delete PCNA at the protein level and achieve superior degradation kinetics over RNAi (RNA interference). Thus, we envision that bioPROTACs can become a valuable research tool for studying the function of long-lived proteins that are known to be refractory to RNAi.

Since PCNA is indispensable for DNA replication^35^, the degradation of PCNA in cells expressing high levels of Con1-SPOP (mCherry^high^) resulted in complete S-phase withdrawal (Fig. 5e top row), as indicated by the lack of cells that stain positive for EdU (a nucleoside analog of thymidine that gets incorporated into newly synthesized DNA). Based on previous reports, the Con1 peptide alone was active as a stoichiometric inhibitor of PCNA since it was able to disrupt the binding of PCNA effector proteins including p21^39,45,46^. Indeed, Con1-SPOP_mut_ (that binds PCNA but was unable to induce its degradation) prevented DNA synthesis but was only effective at high concentrations (Fig. 5e top row). In cells with lower expression of Con1-SPOP_mut_ (mCherry^mid^), effects on the cell cycle were lost even though the PCNA degrader Con1-SPOP continued to show robust inhibition (Fig. 5e middle row). This result highlighted the sub-stoichiometric efficacy of PROTAC strategies and the ability to achieve functional effects at reduced doses compared to conventional ‘occupancy-driven’ inhibitors^11^. The failure to undergo DNA replication also translated into robust growth inhibition in cells where the expression of Con1-SPOP was regulated by doxycycline (Fig. 5f). 1 – 100 ng/ml doxycycline concentrations were effective in maintaining complete growth arrest of HEK293 cells over 10 days, while the lower concentrations reduced proliferation rates compared to the non-binding control Con1_mut_-SPOP. At all doxycycline concentrations tested, complete growth arrest could not be achieved with the stoichiometric inhibitor Con1-SPOP_mut_ even though growth was impaired at the higher concentrations. This result highlights that for certain targets (which may be high in abundance, prone to compensatory feedback mechanisms, or have multiple functions that cannot be inhibited through a single binding site), a degradation approach may be needed to achieve the desired functional outcome.

## DISCUSSION

The advent of small molecule based targeted degradation approaches has ignited a paradigm shift in drug discovery and created unique opportunities to tackle historically intractable targets. To capitalize on this general approach, we are using an orthogonal system termed bioPROTACs, where a polypeptide-based binder to the POI is fused directly to an E3 ubiquitin ligase. Unlike their small molecule counterparts, bioPROTAC discovery is not limited by the ligandability of the POI and the E3. In the present study, we showed that bioPROTACs are highly modular in nature and can readily accommodate changes to either the binder or the E3 ligase. This remarkable flexibility enabled us to tap into the rich diversity offered by the UPS. Indeed, 8 out of 10 different mammalian E3 ligases tested in this study gave significant degradation activities. We also demonstrated a capacity to engage novel substrates (GFP-fusion proteins and PCNA) to determine their degradability through a bioPROTAC approach. Although we were able to generate first-generation active bioPROTACs through rationale design, we anticipate that improved versions (catalytic efficiency, protein half-life, optimal subcellular localization etc.) can be obtained through further protein engineering/maturation and the optimization of binding affinity/specificity, linker length, stability, and E3 selection.

While we are optimistic of the potential of bioPROTACs as a therapeutic modality, they can also serve to enable the development of small molecule based degraders. In particular, bioPROTACs are poised to address key questions that should be understood prior to initiating small molecule based programs, including 1) is the POI a good substrate for poly-ubiquitination and proteasomal degradation (i.e. exposed lysines, structurally disordered segment that initiates unfolding at the 26S proteasome^33^), 2) which E3 ligases are the most effective at inducing its degradation (correct cellular expression of the entire ubiquitination machinery, prolific activity under the relevant diseased state, etc), and 3) what are the functional consequences associated with its degradation. Of significant advantage is the ability of the bioPROTAC approach to rapidly provide these insights to evaluate the feasibility of a small molecule based campaign before embarking on resource-intensive medicinal chemistry.

For target validation purposes, bioPROTACs offer a novel way to deplete a target at the protein level – one that is complementary but offers distinct benefits to the more established RNAi and CRISPR approaches (Fig. 6). Key advantages of bioPROTAC-mediated silencing include 1) insights can be gained quickly as knockdown is achieved over a shorter time period and is not dependent on the natural turnover of the existing pool of proteins (RNAi) nor does it require genetic manipulation (CRISPR), 2) specificity is defined by the substrate binder which can be optimized through yeast- and phage-display technologies, 3) specific post-translational modifications (eg. phosphorylation) or protein state (eg. aggregated) can be selectively targeted, 4) effect of knockdown is reversible – once the bioPROTAC is gone, new proteins can be made to replenish the deleted pool, 5) knockdown can be controlled temporally and/or spatially in animal models through genetic engineering. For example, knock-in mice where the bioPROTAC expression is under the control of a doxycycline-inducible promoter combined with a tissue-specific Tet regulator protein will allow one to delete the bioPROTAC’s substrate in a specific tissue at a specific time in the mouse development. The substrate can also be re-accumulated by withdrawing doxycycline and discontinuing the expression of the bioPROTAC. With the continual advancements in yeast- and phage-display technologies, it is not difficult to identify a peptide or a mini-protein binder against a POI. We showed that as long as the binding affinity is high, it can be combined with the wealth of E3 ligases to generate active bioPROTACs with robust silencing activity. This ease of development should encourage its broad utility as a general proteome editing tool. Moreover, unlike small molecule PROTAC, bioPROTACs can be integrated into the genome making the precise modulation of protein levels for the study of protein function possible.

**Figure 6.**
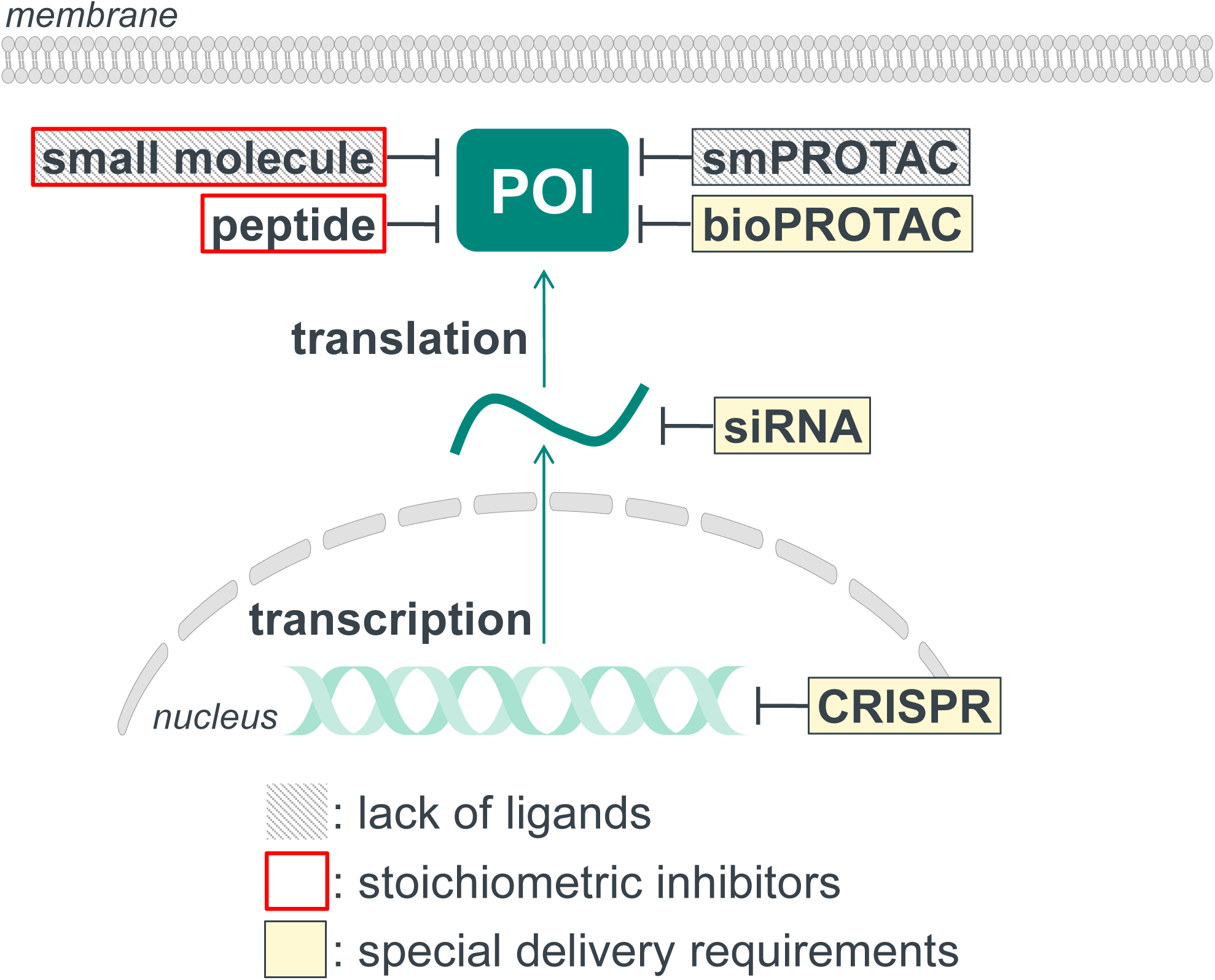

We have also put forth a systematic workflow to guide the selection of binder-E3 ligase pairs. As an illustrative example, we generated a highly effective bioPROTAC against an endogenous protein PCNA, which can be used as a research tool and/or potentially as a cancer therapeutic. The workflow started with the testing of our library of vhhGFP4-based anti-GFP bioPROTACs as degraders of PCNA-GFP. The advantage of this approach is that early insights with respect to a target’s degradability can be gained since a specific binder to the POI is not required at this stage. Indeed, we observed significant degradation of the PCNA-GFP fusion protein with 5 out of 10 vhhGFP4-E3 combinations (Fig. 4). Since GFP itself was a poor substrate for poly-ubiquitin mediated degradation with these constructs (Fig. 4), PCNA is likely amenable for targeted degradation strategies. We were thus motivated to engineer bioPROTACs against endogenous PCNA. This was accomplished by leveraging Con1, a high affinity peptide ligand of PCNA. Indeed, Con1-SPOP proved to give rapid and robust degradation of endogenous PCNA. More importantly, we achieved the expected functional effects in terms of cell cycle arrest and inhibition of cellular proliferation with the active bioPROTAC but not the non-binder control, indicating that on-target biological effects could be seen with the potential to advance this molecule towards the clinic.

To realize the potential of bioPROTACs as a novel therapeutic modality, the fundamental challenge of delivery will need to be addressed. With this in mind, delivery of mRNAs encoding for a bioPROTAC seems particularly attractive as human clinical proof of concept has been achieved with several RNA candidates presently in late stage clinical trials^47^. The optimization of the RNA delivery vehicle as well as the modification of RNA to increase stability and reduce immune activation were key drivers of these latest developments. We anticipate that this will be a viable route for the intracellular delivery of bioPROTACs encoded by mRNA.

During the final stages of manuscript preparation, an orthogonal study by the DeLisa group was published. Although that work focused on leveraging IpaH9.8, a bacterial E3 mimic, for achieving targeted degradation, some interesting parallels were noted. First, using IpaH9.8 as the E3 of choice, a variety of GFP- and GFP-derived-fusion proteins could be degraded. In addition, they also noted some flexibility in terms of the substrate binder that could be used. Together with the work presented here, the combined results reinforce the modular nature of the bioPROTAC approach and one that can be used as a novel biological tool and potential therapeutic.

## ACKNOWLEDGEMENTS

We thank Tomi K. Sawyer, Chandra Verma, Srinivasaraghavan Kannan, Tsz Ying Yuen, Cynthia R. Coffill, Farid J. Ghadessy and all members of the TMRC Pharmacology team for helpful discussions and comments on the manuscript.

## MATERIALS AND METHODS

### Plasmids

To generate Tet-On® 3G bidirectional inducible plasmids, gBlocks® gene fragments for the following inserts were synthesized by Integrated DNA technologies (IDT) and cloned into pTRE3G-BI-mCherry (Clontech) using BamHI and NotI: NSlmb_1-198_-vhhGFP4-FLAG, FLAG-vhhGFP4-SPOP_167-374_, FLAG-vhhGFP4mut-SPOP_167-374_, FLAG-vhhGFP4-SPOPmut, FLAG-vhhGFP4mut-SPOPmut, FLAG-3G86.32-SPOP_167-374_, FLAG-bGFP-A-SPOP_167-374_, FLAG-bGFP-C-SPOP_167-374_, FLAG-GS2-SPOP_167-374_, FLAG-GL6-SPOP_167-374_, FLAG-GL8-SPOP_167-374_, FLAG-βTrCP_2-263_-vhhGFP4, FLAG-FBW7_2-293_-vhhGFP4, FLAG-SKP2_2-147_-vhhGFP4, FLAG-vhhGFP4-VHL_152-213_, FLAG-CRBN_2-320_-vhhGFP4, FLAG-DDB2_2-114_-vhhGFP4, FLAG-vhhGFP4-SOCS2_143-198_, FLAG-vhhGFP4-ASB1_266-335_, FLAG-vhhGFP4-CHIP_128-303_, FLAG-Con1-SPOP_167-374_, FLAG-Con1mut-SPOP_167-374_ and FLAG-Con1-SPOPmut. vhhGFP4mut lacks complementarity determining region 3 (ΔNVNVGFE). SPOPmut lacks the 3-box motif responsible for binding to Cullin (ΔAAEILILADLHSADQLKTQAVDFIN). Con1mut contains 3 point mutations (SAVLQKK<u>A</u>TD<u>AA</u>HPKK). gBlocks for GFP, H2B-GFP, PCNA-GFP were synthesized by IDT and cloned into pEF6 (Thermo Fisher Scientific) using BamHI and NotI. FLAG-Con1-SPOP_167-374_, FLAG-Con1mut-SPOP_167-374_ and FLAG-Con1-SPOPmut were subcloned into pcDNA™4/TO (Thermo Fisher Scientific) using BamHI and NotI. All plasmids were verified by sequencing at 1st BASE.

### Cell culture and transfection

HEK 293 Tet-On® 3G cells were purchased from Clontech and cultured in Minimum Essential Medium (MEM) GlutaMAX™ (Gibco) supplemented with 10% Tet system approved FBS (Clontech) and 100 µg/ml geneticin. T-REx™-293 cells were purchased from Thermo Fisher Scientific and cultured in MEM GlutaMAX™ supplemented with 10% Tet system approved FBS (Clontech) and 5 µg/ml blasticidin. Cells were seeded in poly-D-lysine coated plates and transfected with FuGENE® HD (Promega) the following day according to the manufacturer’s protocol. To induce expression from the pTRE3G-BI-mCherry plasmids, 100 ng/ml doxycycline (Clontech) was added 24 hours post-transfection. To generate stable cell lines expressing GFP, H2B-GFP or PCNA-GFP, HEK 293 Tet-On® 3G cells were selected using 10 µg/ml blasticidin 3 days post-transfection and maintained in 5 µg/ml blasticidin once stable colonies are formed. GFP-positive cells were subsequently enriched by fluorescence-activated cell sorting (FACs) on BD FACSAria™ Fusion. To generate stable cell lines expressing FLAG-Con1-SPOP_167-374_, FLAG-Con1mut-SPOP_167-374_ or FLAG-Con1-SPOPmut, T-REx™-293 cells were selected using 400 µg/ml Zeocin 3 days post-transfection and maintained in 200 µg/ml Zeocin once stable colonies are formed. All cells were maintained at 37ºC, 5% CO_2_ and 90% relative humidity.

### Flow cytometric analysis, cell sorting and EdU labeling

Cells were seeded in 24-well poly-D-lysine coated plates and transfected as described above. 24 hours after transfection, the transfection media was removed and replaced with fresh media containing 100 ng/ml doxycycline. 24 hours after dox-induction, cells were trypsinized and resuspended in cold PBS containing 10% FBS. The cell suspension was passed through a 35 µm nylon mesh to dissociate aggregates before analysis on BD LSRFortessa™ X-20. To sort cells according to mCherry or GFP expression, cells were seeded in 60 mm dishes and harvested in complete media after transfection and dox-induction. A four-way sort was used on BD FACSAria™ Fusion to achieve a purity >98% and a yield >80%. 100, 000 cells were collected and processed for Western blot analysis or for further expansion in culture. For EdU labeling, cells were seeded in 6-well poly-D-lysine coated plates and transfected as described above. Cells were pulse labeled for 2 hours with 10 µM EdU and stained using the Click-iT EdU Flow Cytometry Assay Kit (Thermo Fisher Scientific) according to the manufacturer’s protocol. FxCycle™ Far Red Stain (Thermo Fisher Scientific) was included to determine DNA content. Samples were analyzed on BD FACSAria™ Fusion.

### Imaging

Cells were seeded in 96-well poly-D-lysine coated µCLEAR® plates (Greiner) and allowed to attach overnight. The next day, transfection and dox-induction were performed as described above. Images of live cells were acquired at 37ºC, 5% CO_2_ using the Opera Phenix™ High Content Confocal Screening System under the 20X water immersion lenses. Percentage confluency of cells were tracked continuously over 10 days using the IncuCyte® S3 Live-Cell Analysis System under the 4X whole well imaging objective.

### Western blot analysis

Cells were lysed in ice-cold cell lysis buffer (Cell Signaling Technology) supplemented with 1 mM PMSF and cOmplete™ EDTA-free protease inhibitor cocktail (Roche) for 30 min with intermittent vortexing. Lysates were centrifuged at 18,000 g, 4°C for 15 min and supernatants were snap frozen in liquid nitrogen. Protein concentration was determined using the BCA protein assay kit (Pierce). 20 to 50 μg of protein extract was separated on 4-12% Bis-Tris plus gels, transferred onto nitrocellulose membranes using the Trans-Blot® Turbo™ semi-dry system (Bio-rad), and blocked for 1 hour at room temperature with tris-buffered saline (TBS) Odyssey blocking buffer (Li-Cor). Blots were probed with the appropriate primary antibodies overnight at 4°C in Odyssey blocking buffer supplemented with 0.1% Tween-20, followed by the secondary antibodies IRDye® 680RD donkey anti-mouse IgG and IRDye® 800CW donkey anti-rabbit IgG (Li-Cor) for 1 hour at room temperature. Fluorescent signals were imaged and quantified using Odyssey® CLx. Primary antibodies used were: GFP (Santa Cruz Biotechnology, sc-9996), Hsp90 (BD Transduction Laboratories, 610419), FLAG-tag (Cell Signaling Technology, #8146 and #14793), PCNA (Cell Signaling Technology, #2586 and #13110).

---

**Supplementary Figure 1** Confocal imaging analysis of H2B-GFP/HEK293 Tet-On® 3G cells transiently transfected with various bidirectional, Tet-responsive plasmids. 1 µg/ml doxycycline was added for 24 hours to induce the simultaneous expression of mCherry only or mCherry and NSlmb_1-198_-vhhGFP4-FLAG.

**Supplementary Figure 2** Confocal imaging analysis of HEK293 Tet-On® 3G cells with stable integration of H2B-GFP, GFP, or GFP targeted to different subcellular localizations. Nuclear-targeting of GFP was achieved through the addition of a nuclear localization signal (NLS) from the SV40 large T-antigen. Membrane-targeting of GFP was achieved through the addition of either a palmitoylation signal from Neuromodulin or a farnesylation signal from H-Ras. Images in the top row indicate the correct subcellular distribution of each GFP protein. Each stable cell line was then transiently transfected and induced with doxycycline to drive the co-expression of mCherry and vhhGFP4-SPOP_167-374_.

## REFERENCES

1 Toure, M. & Crews, C. M. Small-Molecule PROTACS: New Approaches to Protein Degradation. Angew Chem Int Ed Engl 55, 1966–1973, doi:10.1002/anie.201507978 (2016).

2 Gechijian, L. N. et al. Functional TRIM24 degrader via conjugation of ineffectual bromodomain and VHL ligands. Nat Chem Biol 14, 405–412, doi:10.1038/s41589-018-0010-y (2018).

3 Huang, X. & Dixit, V. M. Drugging the undruggables: exploring the ubiquitin system for drug development. Cell Res 26, 484–498, doi:10.1038/cr.2016.31 (2016).

4 Churcher, I. Protac-Induced Protein Degradation in Drug Discovery: Breaking the Rules or Just Making New Ones? J Med Chem 61, 444–452, doi:10.1021/acs.jmedchem.7b01272 (2018).

5 Burslem, G. M. et al. The Advantages of Targeted Protein Degradation Over Inhibition: An RTK Case Study. Cell Chem Biol 25, 67–77 e63, doi:10.1016/j.chembiol.2017.09.009 (2018).

6 Bondeson, D. P. et al. Lessons in PROTAC Design from Selective Degradation with a Promiscuous Warhead. Cell Chem Biol 25, 78–87 e75, doi:10.1016/j.chembiol.2017.09.010 (2018).

7 Gadd, M. S. et al. Structural basis of PROTAC cooperative recognition for selective protein degradation. Nat Chem Biol 13, 514–521, doi:10.1038/nchembio.2329 (2017).

8 Huang, H. T. et al. A Chemoproteomic Approach to Query the Degradable Kinome Using a Multi-kinase Degrader. Cell Chem Biol 25, 88–99 e86, doi:10.1016/j.chembiol.2017.10.005 (2018).

9 Smith, B. E. et al. Differential PROTAC substrate specificity dictated by orientation of recruited E3 ligase. Nat Commun 10, 131, doi:10.1038/s41467-018-08027-7 (2019).

10 Collins, I., Wang, H., Caldwell, J. J. & Chopra, R. Chemical approaches to targeted protein degradation through modulation of the ubiquitin-proteasome pathway. Biochem J 474, 1127–1147, doi:10.1042/BCJ20160762 (2017).

11 Lai, A. C. & Crews, C. M. Induced protein degradation: an emerging drug discovery paradigm. Nat Rev Drug Discov 16, 101–114, doi:10.1038/nrd.2016.211 (2017).

12 Edmondson, S. D., Yang, B. & Fallan C. Proteolysis targeting chimeras (PROTACs) in ‘beyond rule-of-five’ chemical space: Recent progress and future challenges. Bioorg Med Chem Lett 29, 1555–1564, doi:10.1016/j.bmcl.2019.04.030 (2019).

13 Cong, F., Zhang, J., Pao, W., Zhou, P. & Varmus, H. A protein knockdown strategy to study the function of beta-catenin in tumorigenesis. BMC Mol Biol 4, 10, doi:10.1186/1471-2199-4-10 (2003).

14 Su, Y., Ishikawa, S., Kojima, M. & Liu, B. Eradication of pathogenic beta-catenin by Skp1/Cullin/F box ubiquitination machinery. Proc Natl Acad Sci U S A 100, 12729–12734, doi:10.1073/pnas.2133261100 (2003).

15 Liu, J., Stevens, J., Matsunami, N. & White, R. L. Targeted degradation of beta-catenin by chimeric F-box fusion proteins. Biochem Biophys Res Commun 313, 1023–1029 (2004).

16 Ma, Y. et al. Targeted degradation of KRAS by an engineered ubiquitin ligase suppresses pancreatic cancer cell growth in vitro and in vivo. Mol Cancer Ther 12, 286–294, doi:10.1158/1535-7163.MCT-12-0650 (2013).

17 Hatakeyama, S., Watanabe, M., Fujii, Y. & Nakayama, K. I. Targeted destruction of c-Myc by an engineered ubiquitin ligase suppresses cell transformation and tumor formation. Cancer Res 65, 7874–7879, doi:10.1158/0008-5472.CAN-05-1581 (2005).

18 Chen, W., Lee, J., Cho, S. Y. & Fine, H. A. Proteasome-mediated destruction of the cyclin a/cyclin-dependent kinase 2 complex suppresses tumor cell growth in vitro and in vivo. Cancer Res 64, 3949–3957, doi:10.1158/0008-5472.CAN-03-3906 (2004).

19 Zhou, P., Bogacki, R., McReynolds, L. & Howley, P. M. Harnessing the ubiquitination machinery to target the degradation of specific cellular proteins. Mol Cell 6, 751–756 (2000).

20 Zhang, J., Zheng, N. & Zhou, P. Exploring the functional complexity of cellular proteins by protein knockout. Proc Natl Acad Sci U S A 100, 14127–14132, doi:10.1073/pnas.2233012100 (2003).

21 Portnoff, A. D., Stephens, E. A., Varner, J. D. & DeLisa, M. P. Ubiquibodies, synthetic E3 ubiquitin ligases endowed with unnatural substrate specificity for targeted protein silencing. J Biol Chem 289, 7844–7855, doi:10.1074/jbc.M113.544825 (2014).

22 Caussinus, E., Kanca, O. & Affolter, M. Fluorescent fusion protein knockout mediated by anti-GFP nanobody. Nat Struct Mol Biol 19, 117–121, doi:10.1038/nsmb.2180 (2011).

23 Shin, Y. J. et al. Nanobody-targeted E3-ubiquitin ligase complex degrades nuclear proteins. Sci Rep 5, 14269, doi:10.1038/srep14269 (2015).

24 Fulcher, L. J. et al. An affinity-directed protein missile system for targeted proteolysis. Open Biol 6, doi:10.1098/rsob.160255 (2016).

25 Saerens, D. et al. Identification of a universal VHH framework to graft non-canonical antigen-binding loops of camel single-domain antibodies. J Mol Biol 352, 597–607, doi:10.1016/j.jmb.2005.07.038 (2005).

26 Rothbauer, U. et al. Targeting and tracing antigens in live cells with fluorescent nanobodies. Nat Methods 3, 887–889, doi:10.1038/nmeth953 (2006).

27 Nagai, Y. et al. Identification of a novel nuclear speckle-type protein, SPOP. FEBS Lett 418, 23–26 (1997).

28 Long, M. J., Poganik, J. R. & Aye, Y. On-Demand Targeting: Investigating Biology with Proximity-Directed Chemistry. J Am Chem Soc 138, 3610–3622, doi:10.1021/jacs.5b12608 (2016).

29 Brauchle, M. et al. Protein interference applications in cellular and developmental biology using DARPins that recognize GFP and mCherry. Biol Open 3, 1252–1261, doi:10.1242/bio.201410041 (2014).

30 Chevrel, A. et al. Specific GFP-binding artificial proteins (alphaRep): a new tool for in vitro to live cell applications. Biosci Rep 35, doi:10.1042/BSR20150080 (2015).

31 Koide, A., Wojcik, J., Gilbreth, R. N., Hoey, R. J. & Koide, S. Teaching an old scaffold new tricks: monobodies constructed using alternative surfaces of the FN3 scaffold. J Mol Biol 415, 393–405, doi:10.1016/j.jmb.2011.12.019 (2012).

32 Bosu, D. R. & Kipreos, E. T. Cullin-RING ubiquitin ligases: global regulation and activation cycles. Cell Div 3, 7, doi:10.1186/1747-1028-3-7 (2008).

33 Guharoy, M., Bhowmick, P., Sallam, M. & Tompa, P. Tripartite degrons confer diversity and specificity on regulated protein degradation in the ubiquitin-proteasome system. Nat Commun 7, 10239, doi:10.1038/ncomms10239 (2016).

34 Marzahn, M. R. et al. Higher-order oligomerization promotes localization of SPOP to liquid nuclear speckles. EMBO J 35, 1254–1275, doi:10.15252/embj.201593169 (2016).

35 Moldovan, G. L., Pfander, B. & Jentsch, S. PCNA, the maestro of the replication fork. Cell 129, 665–679, doi:10.1016/j.cell.2007.05.003 (2007).

36 Wang, S. C. PCNA: a silent housekeeper or a potential therapeutic target? Trends Pharmacol Sci 35, 178–186, doi:10.1016/j.tips.2014.02.004 (2014).

37 Chen, J., Bozza, W. & Zhuang, Z. Ubiquitination of PCNA and its essential role in eukaryotic translesion synthesis. Cell Biochem Biophys 60, 47–60, doi:10.1007/s12013-011-9187-3 (2011).

38 Mailand, N., Gibbs-Seymour, I. & Bekker-Jensen, S. Regulation of PCNA-protein interactions for genome stability. Nat Rev Mol Cell Biol 14, 269–282, doi:10.1038/nrm3562 (2013).

39 Zheleva, D. I. et al. A quantitative study of the in vitro binding of the C-terminal domain of p21 to PCNA: affinity, stoichiometry, and thermodynamics. Biochemistry 39, 7388–7397 (2000).

40 Senga, T. et al. PCNA is a cofactor for Cdt1 degradation by CUL4/DDB1-mediated N-terminal ubiquitination. J Biol Chem 281, 6246–6252, doi:10.1074/jbc.M512705200 (2006).

41 Yu, Y. et al. Proliferating cell nuclear antigen is protected from degradation by forming a complex with MutT Homolog2. J Biol Chem 284, 19310–19320, doi:10.1074/jbc.M109.015289 (2009).

42 Xu, B. et al. Proliferating cell nuclear antigen (PCNA) regulates primordial follicle assembly by promoting apoptosis of oocytes in fetal and neonatal mouse ovaries. PLoS One 6, e16046, doi:10.1371/journal.pone.0016046 (2011).

43 Niimi, A. et al. Regulation of proliferating cell nuclear antigen ubiquitination in mammalian cells. Proc Natl Acad Sci U S A 105, 16125–16130, doi:10.1073/pnas.0802727105 (2008).

44 Schwanhausser, B. et al. Global quantification of mammalian gene expression control. Nature 473, 337–342, doi:10.1038/nature10098 (2011).

45 Kontopidis, G. et al. Structural and biochemical studies of human proliferating cell nuclear antigen complexes provide a rationale for cyclin association and inhibitor design. Proc Natl Acad Sci US A 102, 1871–1876, doi:10.1073/pnas.0406540102 (2005).

46 Warbrick, E. A functional analysis of PCNA-binding peptides derived from protein sequence, interaction screening and rational design. Oncogene 25, 2850–2859, doi:10.1038/sj.onc.1209320 (2006).

47 Kaczmarek, J. C., Kowalski, P. S. & Anderson, D. G. Advances in the delivery of RNA therapeutics: from concept to clinical reality. Genome Med 9, 60, doi:10.1186/s13073-017-0450-0 (2017).

