## Supplementary figures and images for "bioPROTACs as versatile modulators of intracellular therapeutic targets: Application to proliferating cell nuclear antigen (PCNA)" (from another version of this preprint: version 1, 2019-08-07)

### Supplementary figures

#### Supplementary Figure 1

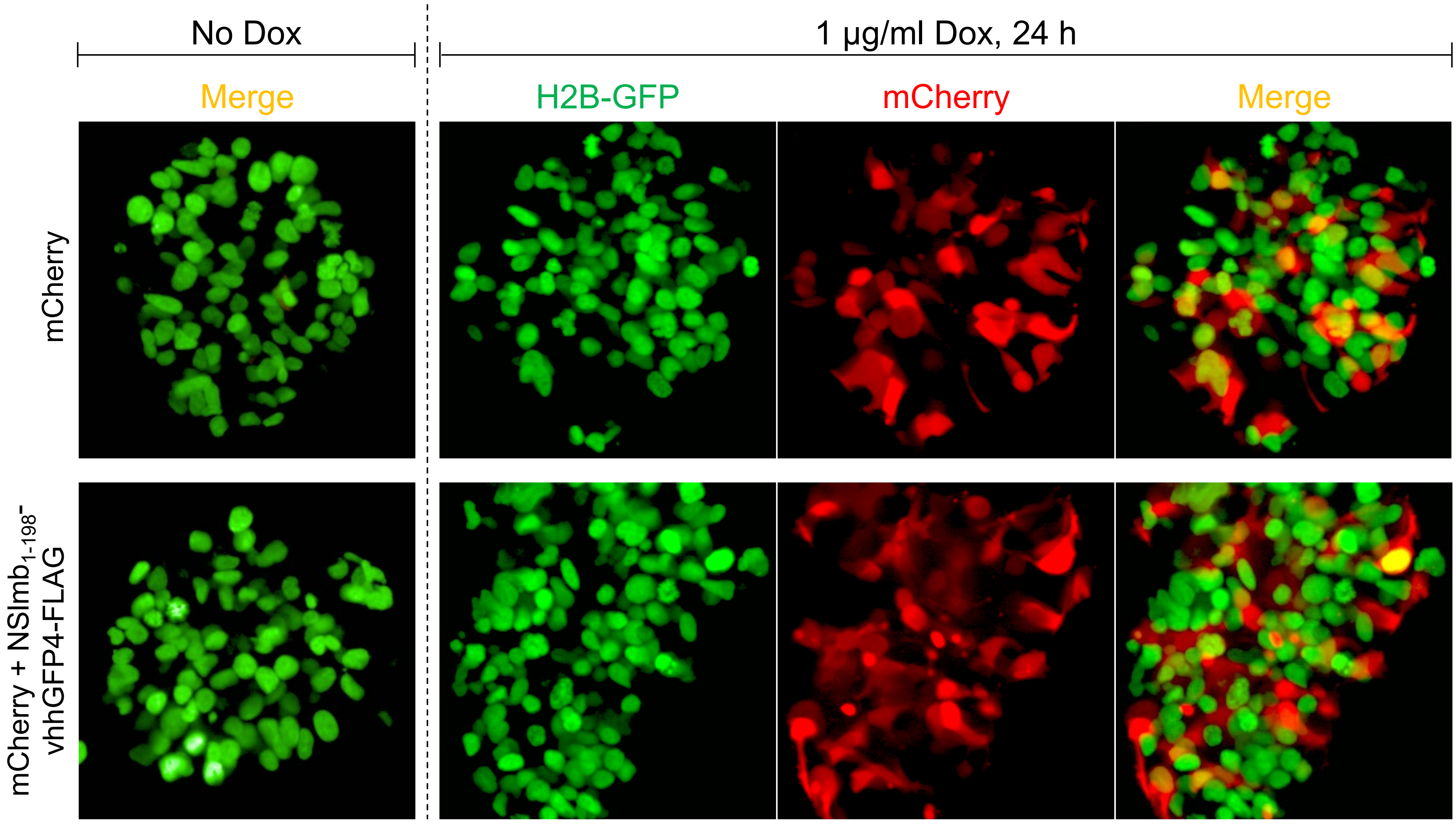


Public

#### Supplementary Figure 2

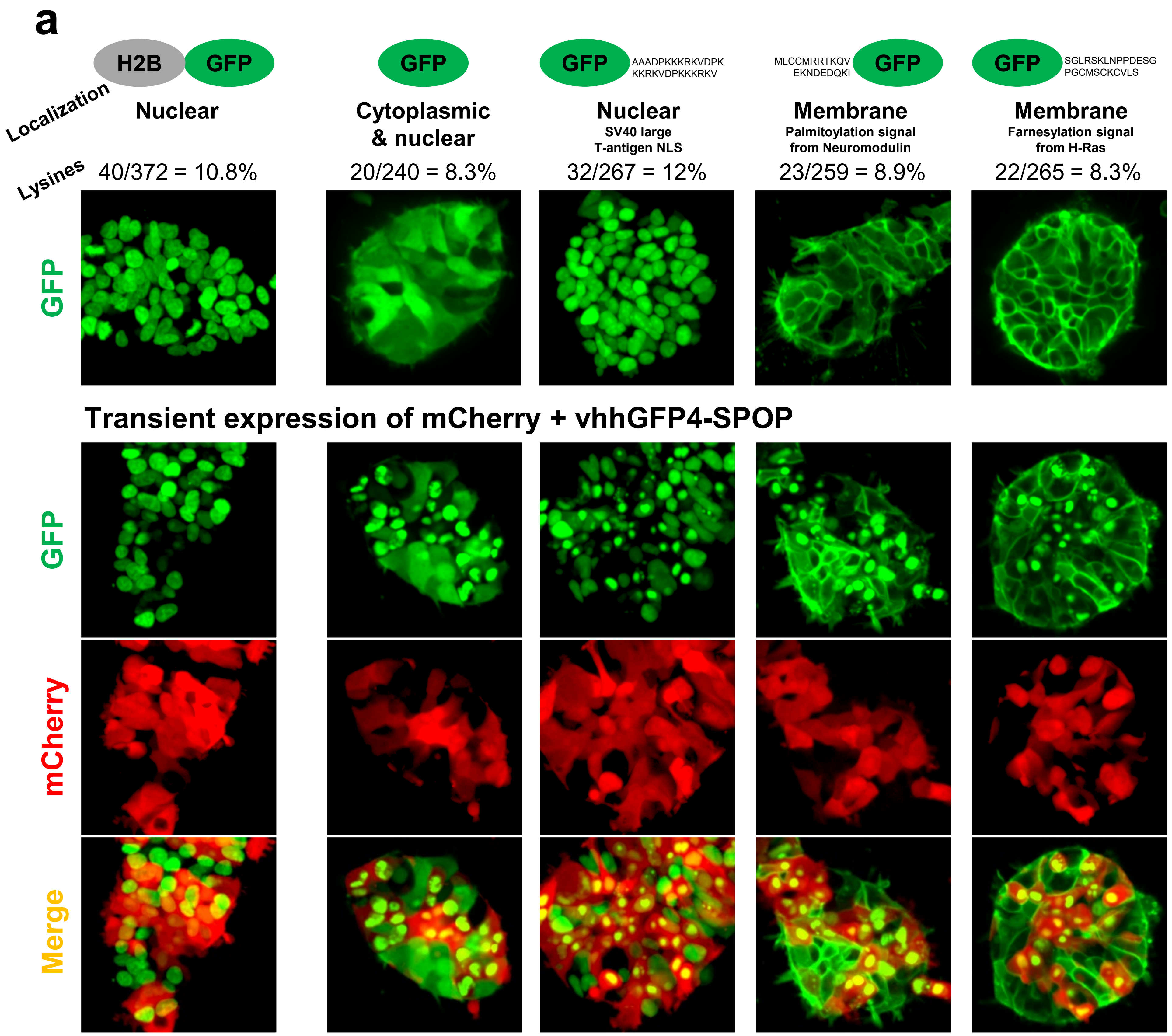


Public
